# Replication Timing Networks: a novel class of gene regulatory networks

**DOI:** 10.1101/186866

**Authors:** Juan Carlos Rivera-Mulia, Sebo Kim, Haitham Gabr, Abhijit Chakraborty, Ferhat Ay, Tamer Kahveci, David M. Gilbert

**Author notes:** Contact Information: David M. Gilbert, Department of Biological Science, Florida State University 319 Stadium Drive, Tallahassee, FL 32306-4295.

## Abstract

DNA replication occurs in a defined temporal order known as the replication-timing (RT) program and is regulated during development, coordinated with 3D genome organization and transcriptional activity. However, transcription and RT are not sufficiently coordinated to predict each other, suggesting an indirect relationship. Here, we exploit genome-wide RT profiles from 15 human cell types and intermediate differentiation stages derived from human embryonic stem cells to construct different types of RT regulatory networks. First, we constructed networks based on the coordinated RT changes during cell fate commitment to create highly complex RT networks composed of thousands of interactions that form specific functional sub-network communities. We also constructed directional regulatory networks based on the order of RT changes within cell lineages and identified master regulators of differentiation pathways. Finally, we explored relationships between RT networks and transcriptional regulatory networks (TRNs), by combining them into more complex circuitries of composite and bipartite networks, revealing novel *trans* interactions between transcription factors and downstream RT changes that were validated with ChIP-seq data. Our findings suggest a regulatory link between the establishment of cell type specific TRNs and RT control during lineage specification.

## Introduction

During development, specific transcriptional programs and epigenetic landscapes are established that maintain the identities and functionality of the specialized cell types that emerge. Despite characterization of changes in transcriptome and epigenome during development (Gifford et al. 2013; Xie et al. 2013; Roadmap Epigenomics Consortium et al. 2015; Tsankov et al. 2015), little is known about the role of spatio-temporal genome organization in cell fate specification. Changes in gene activity and chromatin 3D organization are coordinated with dynamic changes in the temporal order of genome duplication, known as the replication timing (RT) program (Hiratani et al. 2010; Rivera-Mulia et al. 2015; Hiratani et al. 2008; Rivera-Mulia et al. 2018a). Spatio-temporal control of RT is conserved in all eukaryotes (Solovei et al. 2016; Rivera-Mulia and Gilbert 2016a; Zhao et al. 2017) and alterations in the RT program are associated with different diseases (Ryba et al. 2012; Gerhardt et al. 2014; Rivera-Mulia et al. 2017; Sasaki et al. 2017). RT is regulated during development in discrete chromosome units, referred to as replication domains (RDs), that align with topological associated domains (TADs) mapped by chromosome conformation capture techniques (Hi-C) and segregate into distinct nuclear compartments visualized by either cytogenetic or Hi-C methods (Jackson and Pombo 1998; Moindrot et al. 2012; Pope et al. 2014; Rivera-Mulia and Gilbert 2016b; Rivera-Mulia et al. 2018a; Ryba et al. 2010; Sadoni et al. 2004; Yaffe et al. 2010). Hence, we reasoned that RT can be exploited to characterize regulatory relationships between 3D genome organization and gene expression control during development.

Previously, we generated the most comprehensive database of RT programs during human lineage specification and found that approximately half of the genome undergoes dynamic changes that are closely coordinated with the establishment of transcriptional programs (Rivera-Mulia et al. 2015). Additionally, we demonstrated that genes within developmentally RT regulated domains are higher in the hierarchy of transcriptional regulatory networks (TRNs) when compared to RT constitutive genes (Rivera-Mulia et al. 2015). However, strong gene expression was not restricted to early replicating genomic regions and transcriptional activation during cell fate commitment often preceded RT changes (Rivera-Mulia et al. 2015; Rivera-Mulia and Gilbert 2016b). In fact, although a long-standing correlation between early replication and gene expression has been observed in all eukaryotes, the link between RT and transcriptional activity is complex and causal relationships have not been established (Rivera-Mulia and Gilbert 2016b; Solovei et al. 2016). Here, we explored the possibility that RT can be regulated by the establishment of complex regulatory circuits of transcription factors rather than by the transcription levels of genes within each RD. Using the RT program for all RefSeq genes in the human genome we constructed distinct types of RT regulatory network models based on: 1) correlation patterns in RT changes during cell fate commitment, 2) the temporal order of RT changes in each developmental transition and 3) combined networks that explore the crosstalk between RT and transcriptional regulatory networks (composite and bipartite networks). Integrating RT and transcriptional regulatory interactions we found a regulatory link between TRNs and RT control during cell differentiation.

## Results

### Construction of correlation-based RT networks

To determine whether the RT program can be used for novel gene regulatory interaction identification, we defined a model that describes the relationship between all possible combinations of gene pairs (nodes), establishing gene interactions (edges) according to their correlated RT patterns during differentiation towards cell types representing the three main germ layers (Figure 1A). Distinct filters were applied in our model: a) we included only genes that change RT during cell differentiation (removing RT constitutive genes; b) we included edges only between genes separated by at least 500kb and/or in different chromosomes (Figure 1B-C and Supplemental Figure S1). Separation by >500kb was chosen to remove gene pairs within the same RD, which we have shown vary in size from 0.4 to 0.8 Mb (Hiratani et al. 2008; Rivera-Mulia et al. 2015; Pope et al. 2014), and include only distal gene interactions; c) we established gene interactions (edges) between significantly correlated gene pairs (statistical significance of gene pair interactions was calculated as Bonferroni’s adjusted *p-values* –see Methods). After applying these filters, gene pairs and interaction edges were obtained (green boxes in Figure 1C) and RT networks constructed. Figure 1D and E illustrate hypothetical examples of two distinct RT patterns along a particular cell differentiation lineage, the correlations for which constitute connections between gene pairs exploited to construct the corresponding RT networks. These findings revealed co-regulated RT changes for thousands of gene pairs during differentiation that are likely to be mediated by common mechanisms related to differentiation. These gene pairs were then used to construct the following networks.

**Figure 1.**
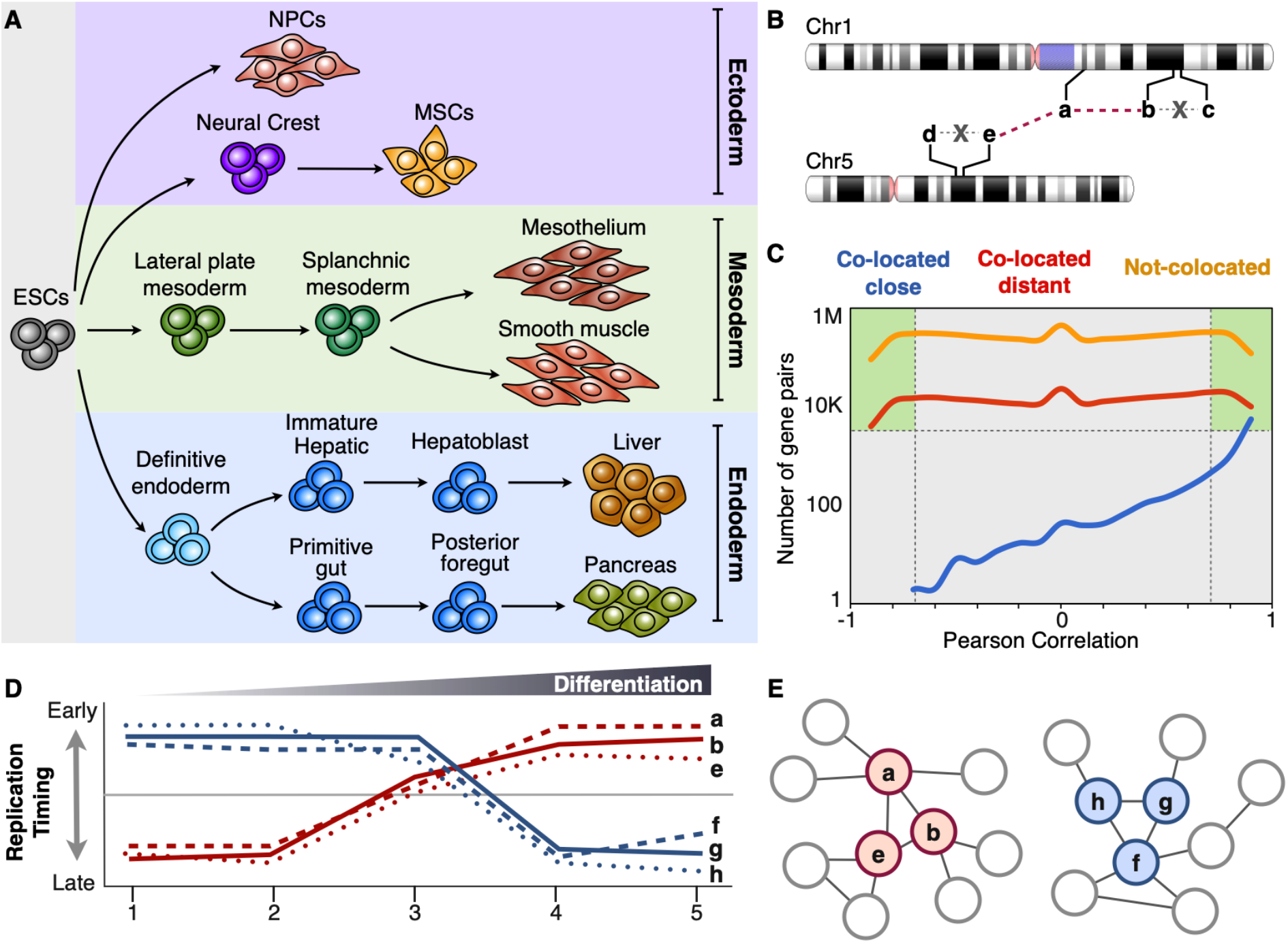
Coordinated changes in RT can be exploited to construct gene regulatory networks (click for full resolution). A) RT programs of distinct cell types representing intermediate stages of human embryonic stem cell differentiation towards endoderm, mesoderm, ectoderm were analyzed for the construction of RT regulatory networks. B) Depiction of different highly RT correlated genes from distinct chromosomes and the establishment of network interaction edges between them. From all possible combinations of gene pairs, those co-located within 500kb were removed from the analysis to include exclusively distal gene interactions. Regulatory interactions (edges) between gene pairs are considered only for genes located >500kb apart (colocated distant) or in distinct chromosomes (not co-located), i.e. edges between genes *b-c* and *d-e* were not included in the analysis. C) Number of gene pairs as function of RT correlation for distinct categories of gene pairs: co-located close (within 500kb), co-located distant (separated by > 500kb) and not co-located (from different chromosomes). Only gene pairs with significant RT correlations and located at least 500kb apart were considered. D) RT patterns during cell fate commitment of distinct hypothetical genes. E) Construction of RT regulatory networks based on the significant *Pearson’s* correlation (distance between nodes are proportional to the correlation strength).

### RT sub-network communities are composed of genes with specific functional annotations

To examine whether co-regulated RT changes are linked to functional gene regulatory interactions, we constructed RT networks among significantly correlated gene pairs and characterized their connectivity and association to specific functional annotations. First, we constructed a RT network including all significantly RT correlated gene-pairs across all differentiation pathways, positioning each node in a two-dimensional space using the force-directed layout algorithm in Cytoscape (Shannon et al. 2003), with the edge length being proportional to the Pearson’s correlation strength (Figure 2A). The resulting RT network was composed of highly interconnected groups of nodes (with >90% of the nodes having a degree distribution >25), with very short distances (path length <3) and high clustering coefficients within each group (Figure 2A). Next, we identified local neighborhoods (sub-network communities) within the global networks according to the connectivity and distances between nodes (Blondel et al. 2008). Finally, to validate the biological significance of the novel gene interactions detected within RT networks, we examined their functional organization by performing ontology analysis of each sub-network community using the spatial analysis of functional enrichment (SAFE) algorithm (Baryshnikova 2016; Costanzo et al. 2016). We found subnetwork communities with highly interconnected nodes of genes involved in specific functions, which were color coded based on the enrichment of functional ontology annotations (Figure 2A). Closer inspection of sub-network communities annotated with specific functions grouped the genes according to their ontology terms (Figure 2B). These findings confirm that novel gene regulatory interactions can be identified exploiting the cell type-specific RT program.

We also generated RT networks of differentiation towards each germ layer separately by combining the cell types of each lineage (ectoderm, mesoderm and endoderm). Then we visualized these RT networks as 2D maps and identified sub-network communities as described above. Consistently, we found RT networks composed by highly interconnected sub-network communities associated with specific functions relevant to each germ layer (Figure 2C-E). Significant enrichment of ontology terms for specific sub-network communities validate the functional relationships between the nodes of these novel interactions. Additional sub-network communities without enrichment in ontology identify co-regulated relationships that have not been previously annotated. Additionally, correlated RT networks vary greatly in size depending on the parameters selected; here we considered only the most significant correlated nodes that switch RT between the very early (>0.3) and very late (<0.3) replication and with > 20-degree count (number of edges). Relaxing the parameters (for early and late replication, significance threshold and degree count) would produce larger networks with higher number of local neighborhoods (at the expense of computational time). Overall, our results revealed that dynamic changes in RT can be exploited to characterize the complex regulatory interactions established during cell fate commitment.

**Figure 2.**
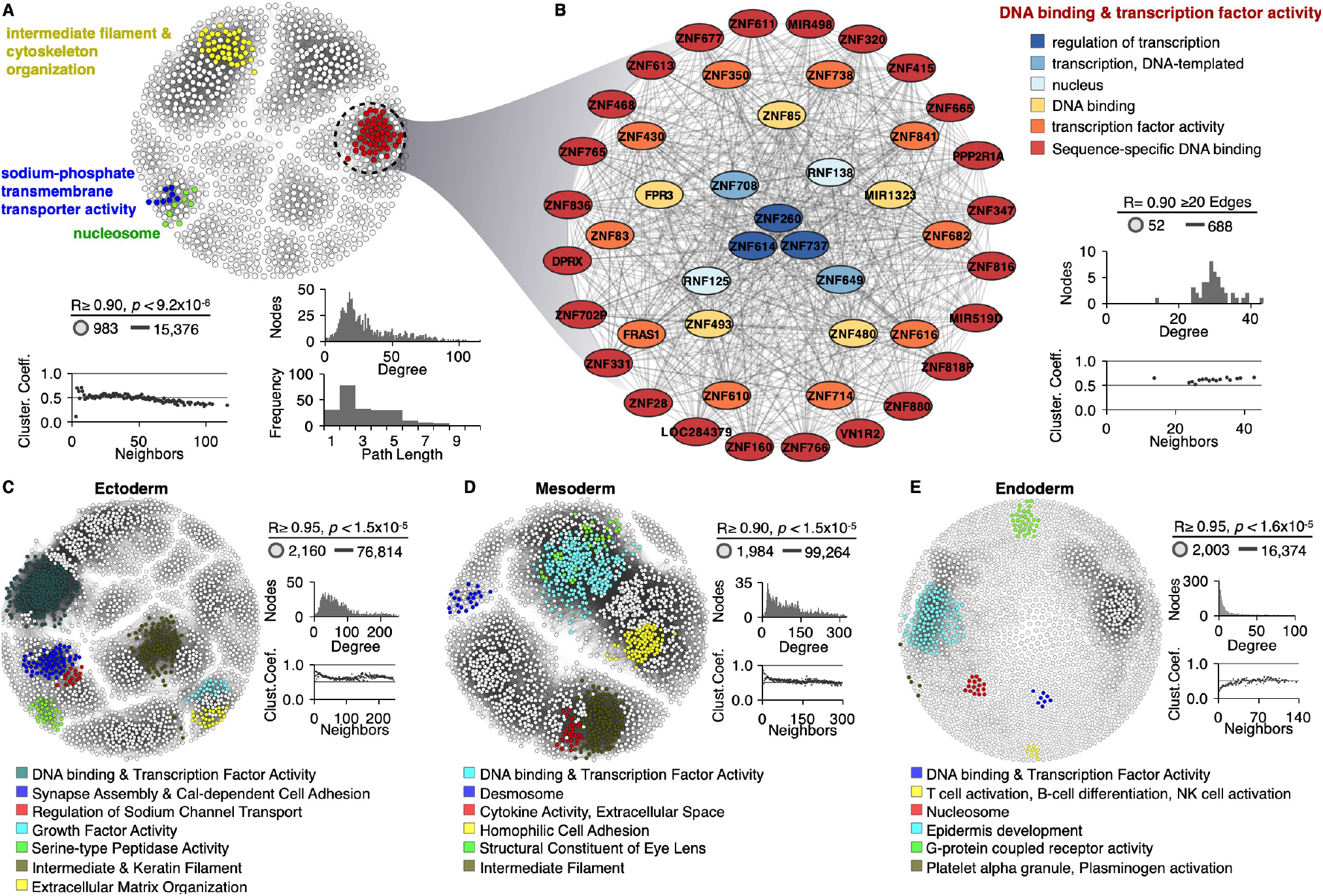
Functional annotation of RT networks (click for full resolution). A) RT co-regulated gene-pairs across all differentiation pathways from human ES cells were identified and exploited to construct a RT network. B) Detailed sub-network connectivity from the RT network shown in A) and its respective node organization per ontology term. C-E) RT networks and functional sub-network communities constructed for each differentiation pathway: ectoderm (C), mesoderm (D) and endoderm (E). Interaction edges between gene pairs were established only for significantly correlated nodes (Bonferroni’s adjusted *p-values* with alpha=0.05/n) and the subset of most connected nodes (>20 edges) were used to visualize RT networks displayed as 2D maps in Cytoscape (Shannon et al. 2003). Pearson’s correlation and Bonferroni’s corrected *p-values* thresholds, as well as the connectivity analysis for each network are shown (degree distribution, path lengths and clustering coefficients). Highly interconnected sub-network communities were annotated with functional ontology terms using SAFE algorithm (Baryshnikova 2016) and displayed in distinct colors.

### Directional RT networks identify regulatory interactions of cell fate commitment

The RT networks described above (Figure 2) identified sub-network communities enriched for specific functional annotations. However, these correlated RT networks are not directional and as such cannot characterize the hierarchical relationships or identify potential targets for regulators. Hence, here we analyzed whether RT can be also exploited to characterize the hierarchical gene regulatory interactions during lineage commitment. To do so, we took advantage of the data collected at distinct intermediate differentiation stages towards pancreas, liver, smooth muscle, mesothelium, mesenchymal stem cells (MSCs) and neural precursors (NPCs) and constructed directional RT networks for each of the specific differentiation pathways. First, we classified the genes according to the order of RT changes during each differentiation pathway (Figure 3A), identified those that change during the earliest cell fate transition, assigned directional edges to genes that changed RT in the subsequent differentiation stage and repeated this for each stage (see Methods). Then, directional RT networks were displayed either in 2D maps or in a hierarchical arrangement and nodes were color/size coded according to the order of the changes in RT during distinct differentiation pathways (Figure 3B). Construction of directional RT networks constitutes a novel approach to identify the genes with earliest RT changes during cell fate commitment (red nodes in Figure 3C) as well as their downstream relationships in terms of temporal ordering of RT change (green, blue and grey nodes in Figure 3C). To test this approach and its value for identifying novel gene regulatory interactions, we constructed directed RT networks for known transcription factors that are key regulators for each differentiation pathway. Consistently, known downstream targets were identified among the downstream targets predicted by the directed RT networks (Supplemental Figure S2). Hence, we classified the nodes in hierarchy levels according to the order of RT changes: “master regulators” were defined as the genes that change RT in the earliest differentiation transition with the largest degree of connectivity (red node in Figure 3C), while downstream nodes were classified according to the time during differentiation when they change RT as managers or effectors (green and blue nodes and grey nodes respectively in Figure 3C). Establishment of RT networks occurs differently for each germ layer, for endoderm cell types (liver and pancreas) most of the changes occur very early during differentiation, with many master regulators changing to early replication and fewer downstream nodes in each of the subsequent differentiation stages (Figure 3D). In contrast, for mesoderm cell types (smooth muscle and mesothelium) fewer master regulators were connected with an increased number of downstream nodes.

The presence of known targets for key differentiation regulators between the predicted downstream targets in the directed RT networks suggest that the interactions identified by this approach reflect functional gene relationships. However, to further validate the gene regulatory interactions within the directional RT networks we constructed directional networks for transcription factors key for the differentiation control towards liver and pancreas. Moreover, during the preparation of this manuscript, ChIP-seq datasets for transcription factors binding became available for liver and pancreas (Diaferia et al. 2016; Wang et al. 2018; https://www.encodeproject.org/), permitting us to validate these downstream targets predicted by the novel RT networks. We constructed directed RT networks for liver and pancreatic differentiation using *FOXA2* and *FOXA1* as source nodes respectively (Figure 3E-F), and found an enrichment for genes with ChIP-seq peaks at their promoters among the predicted downstream targets using RT (Supplemental Table S1). Exemplary target genes and ChIP-seq signals for downstream nodes of the *FOXA2* and *FOXA1* directed RT networks are shown in Figure 3E-F. These findings validate the ability of the directed RT networks to identify novel directional regulatory mechanisms and place them into their temporal order during cell fate commitment.

**Figure 3:**
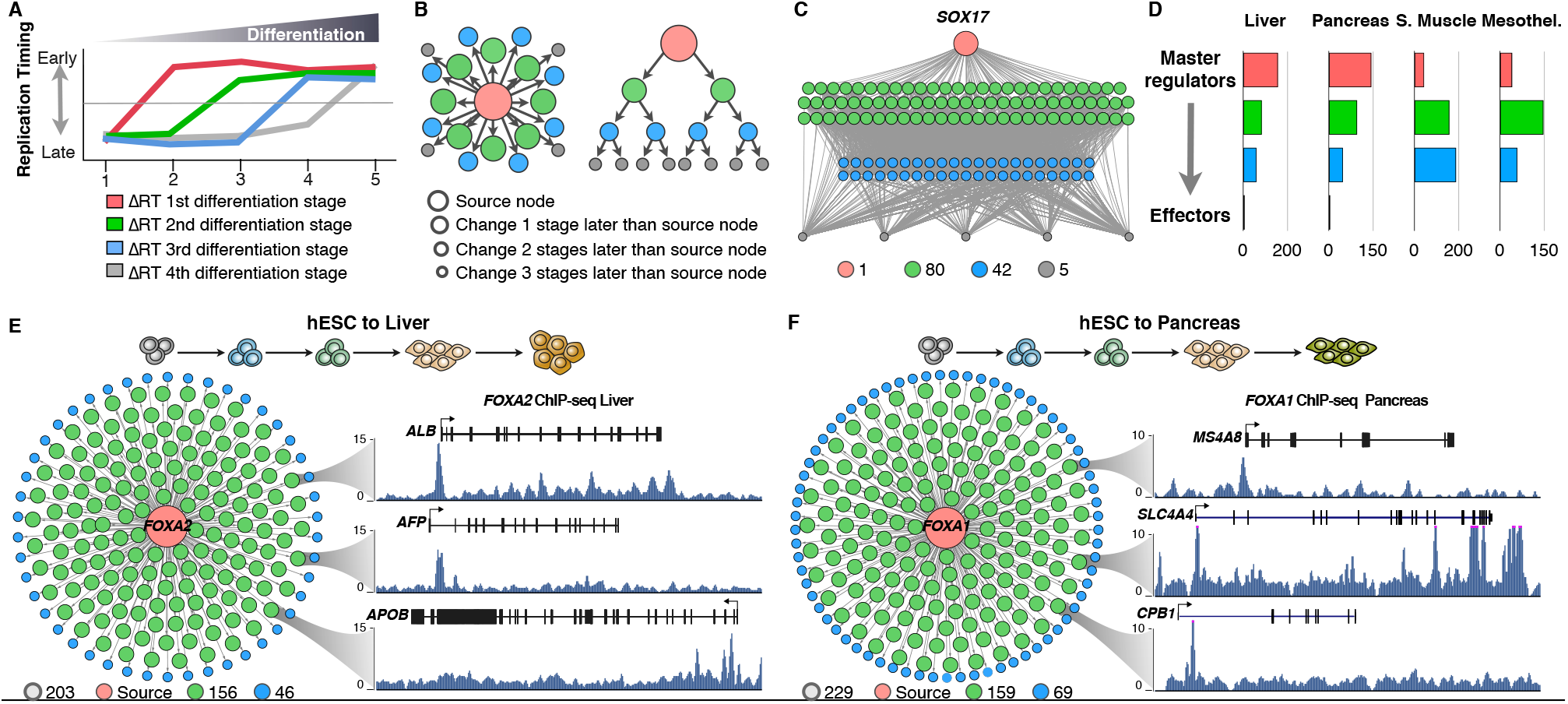
Directional RT networks (click for full resolution). A) Distinct genes change RT at different time points during cell fate commitment and the order of RT changes can be used to construct directional RT networks. Red genes change during the first transition between differentiation stages while grey genes change at the last differentiation transition. B) 2D maps and hierarchical displays of a directional RT networks were constructed based on a source gene (central node) and downstream connected nodes. C) An exemplary directional RT network for liver differentiation is shown. The central node is *SOX17* and all downstream nodes were connected based on temporal times during differentiation at which they change RT. D) Classification of gene nodes according to their hierarchy in directed RT networks. *Master regulators* were operationally defined as those genes that change RT in the earliest differentiation transition and have the largest degree of connectivity (red nodes). Green and blue nodes represent “manager” nodes that are connected to the final “effector” nodes at the lowest level of the network (grey nodes). Node distribution in each of the hierarchical levels for each differentiation pathway is shown. E-F Directed RT networks of *FOXA2* and *FOXA1* during liver and pancreatic differentiation, respectively. *FOXA2* and *FOXA1* (red node) change from late to early replication during the earliest stages of differentiation, potential downstream genes were identified as those genes that change in subsequent differentiation stages (green and blue nodes). ChIP-seq signals show the binding of *FOXA2* and *FOXA1* at the promoters of predicted hepatic-specific and pancreas-specific downstream genes (*ALB, AFP* and *APOB* for liver and *CPB1, MS4A8* and *SLC4A4* for pancreas).

### RT network edges overlap with known transcriptional regulatory interactions

To determine the extent to which RT networks capture the regulatory interactions characterized by other methods, we analyzed their overlap with transcriptional regulatory networks (TRNs) using a previously described set of cell type-specific networks of TFs (Neph et al. 2012). First, we identified the cell types that most closely match the TRNs to our RT networks, as follows: hESC-derived hepatocytes were compared to TRNs from HepG2–a liver cancer cell line that retains morphological and functional hepatocyte properties (Berger et al. 2015; Knowles et al. 1980), hESC-derived mesothelial cells were compared to TRNs from HCF cells –cardiac fibroblasts that during development and *in vitro* differentiation can be derived from mesothelial cells (Mutsaers 2004) and hESC-derived neural precursors were compared to TRNs from the SK-N-SH cell line after treatment with retinoic acid–SK-N-SH cells were derived from a neuroblastoma and retinoic acid causes differentiation to neural phenotype (Preis et al. 1988). Next, we constructed RT networks using only the subset of genes present in the TRNs (475 TFs) that change RT and are significantly correlated in their RT patterns in each differentiation pathway. Finally, we identified the number of common and unique edges between RT networks and TRNs. We found that in all three cases there was a highly significant overlap when compared to the expected overlap by randomly selecting the same number of edges (Figure 4A). In fact, significant overlap was also observed when all cell types from both RT networks and TRNs were classified per germ layer (Supplemental Table S2) and common edges were identified for ectoderm and mesoderm, even when distinct cell types were used for each germ layer. These results confirm a high overlap between RT and transcriptional networks and further validate the gene regulatory interactions identified using the RT program.

**Figure 4:**
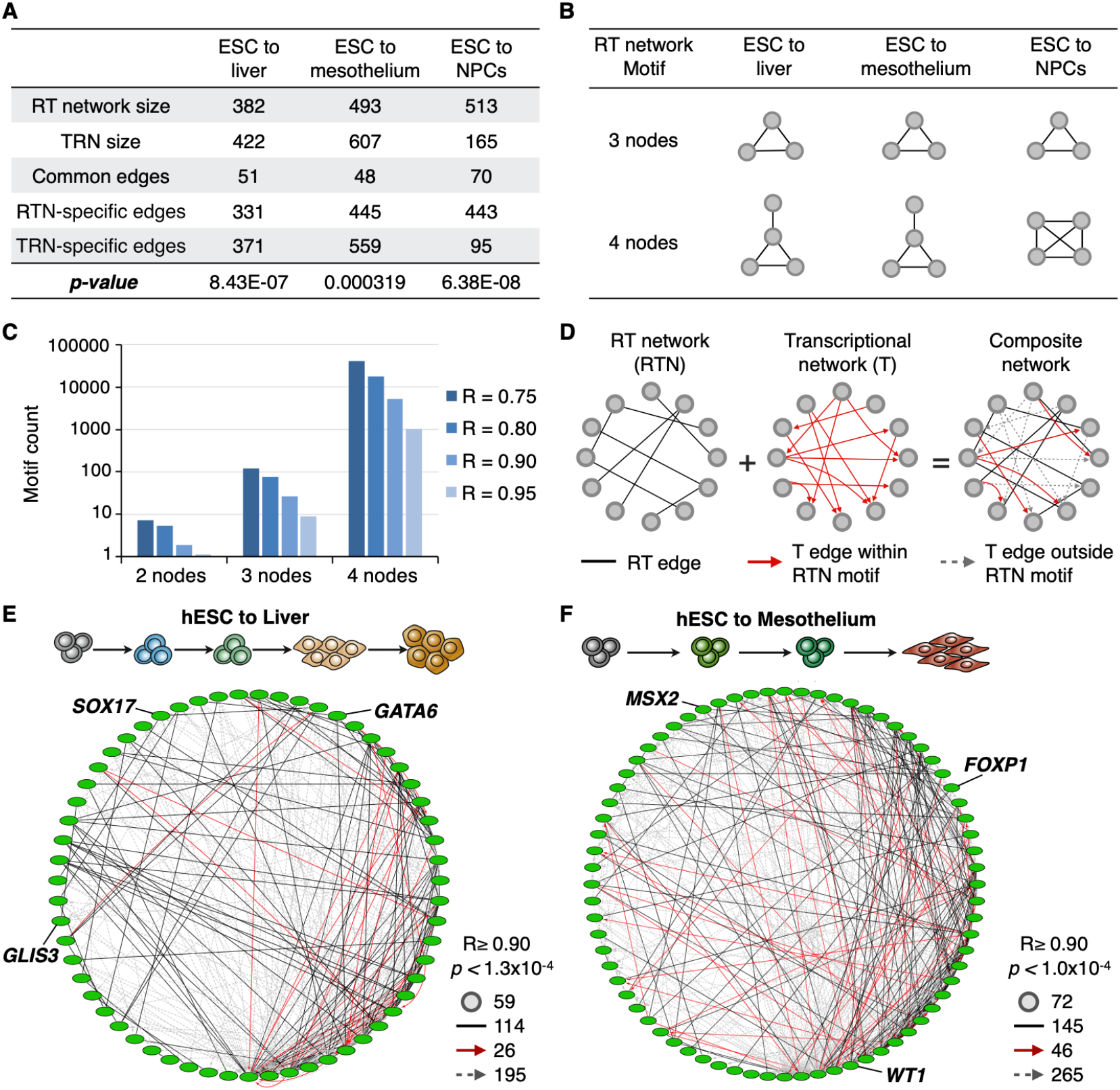
RT and transcriptional networks overlap and can be combined into composite networks (click for full resolution). A) Overlap analysis of RT and TRNs interaction edges. RT networks were constructed for matching cell types in the TRNs (Neph, et al., 2013) and common and unique interaction edges were identified. Only genes within the TRNs were used (475 transcription factors), maintaining the filters for significant correlation and gene pairs *in trans* (separated by >500kb or in distinct chromosomes). Hypergeometric test was performed to test the overlap significance (*p-values* are shown). B-C RT networks are enriched in motifs of ≤ 4 nodes. B) The most enriched motif for each differentiation pathway is shown. C) Motif frequency distribution at distinct thresholds in Pearson’s correlation values. D) Construction of composite networks by combining RT and transcriptional networks. RT networks were used to define the “base network” which include all nodes of RT correlated genes, interaction edges between RT nodes were extracted from TRNs. Composite networks included all RT edges (black undirected lines), transcriptional edges within RT network motifs (directed solid red arrows) and transcriptional edges outside RT network motifs (directed dashed grey arrows). Exemplary composite networks for liver (E) and mesothelium differentiation (F) are shown. Pearson’s correlation and Bonferroni’s corrected *p-values* thresholds, as well as the number of nodes and edges are shown.

### Building blocks of RT networks are motifs with multiple nodes

Previous studies explored the architecture of gene regulatory relationships by analyzing either transcriptional or protein interactions and found that complex cellular networks are constituted by sets of small network motifs, such as interactions between transcription factors and their targets (Alon 2007; Zhang et al. 2005). Here, we performed a topology characterization of RT networks constructed with the subset of genes (475 TFs) present in the TRNs (Neph et al. 2012), to explore the most overrepresented patterns of connectivity. We computed all possible network motifs composed by 2-4 nodes and identified the motifs with high enrichment in each RT network constructed per differentiation pathway (Supplemental Figures S3-S5). Statistical significance (*p-value* <0.01) of each motif occurrence in the RT networks was calculated by comparing to the frequency of the same motif in randomized networks (Milo et al. 2002; Baiser et al. 2015; Elhesha and Kahveci 2016). The most frequent motifs with 2, 3 and 4 nodes in each differentiation pathway are shown in Figure 4B. Previous observations in transcriptional and protein regulatory networks that suggest that gene regulatory networks are composed mainly by small motifs with 2-3 nodes (Yeger-Lotem et al. 2004; Alon 2007). By contrast, we found that the most enriched motifs are the motifs with higher number of nodes (Figure 4C). In fact, motif enrichment increases with higher number of nodes and this distribution is maintained at distinct Pearson’s correlation thresholds and in all differentiation pathways analyzed (Figure 4C). These findings suggest a higher connectivity for RT co-regulated genes than that previously observed in transcriptional and protein regulatory networks. Additionally, since significant overlap between RT and transcriptional networks was observed in all differentiation pathways (Figure 4A), we examined the presence of transcriptional edges within the RT networks motifs. We found that transcriptional edges were present in most of the motifs; however, all frequent motifs contained fewer TRN edges as compared to the RT edges (Supplemental Figures S3–S5).

### Composite networks: combining RT and transcriptional regulatory networks

To better understand how the regulatory circuitries are established during cell fate commitment we constructed a model of composite regulatory networks by merging RT and TRNs. Previous studies demonstrated that distinct types of interactions (such as protein-protein and transcription regulation) can be combined to explore more complex cellular circuitries (Yeger-Lotem et al. 2004; Vidal et al. 2011). Here, we combined the regulatory interactions observed in RT networks with those identified in the TRNs between TFs detected for the closest cell types. First, we used as base networks the set of motifs from each RT network and identified the nodes that are also connected in TRNs (Figure 4C). Then we identified all the interactions between those RT nodes in the TRNs from matching cell types and constructed composite networks by adding the transcriptional interactions (Figure 4C). RT networks for each differentiation pathway are constituted by multiple unconnected motifs of ≤4 nodes; however, the addition of transcriptional edges revealed more complex and highly interconnected networks with all nodes interacting with at least 3 other nodes (Figure 4B-D). Exemplary composite networks for liver and mesothelium show the connectivity of known key regulators for each differentiation pathway (Figure 4E-F).

### Bipartite networks reveal transcription factors as regulators of RT

To further characterize the gene regulatory interactions established during cell fate commitment we analyzed the cell type-specific transcriptomes and its relationships with RT. First, we analyzed genome-wide transcriptomes for the same cell types from which we obtained the RT programs (Rivera-Mulia et al. 2015). Our highly comprehensive characterization of gene expression, including multiple replicates for each differentiation stage, allowed us to identify with confidence the genes that are differentially expressed during cell fate commitment towards each cell type and the genes that better distinguish each intermediate stage. Co-expressed genes were identified by weighted correlation network analysis (Langfelder and Horvath 2008). Strong correlations between gene expression levels are widely used to identify regulatory interactions (Li 2002; Horvath and Dong 2008; D’haeseleer et al. 2000; Gabr et al. 2015; Novak and Jain 2006; Allocco et al. 2004; Laurenti et al. 2013); thus, we used the most significant co-expressed genes to construct TRNs for each differentiation pathway. To decrease the complexity of the data to a computationally manageable size, we focused on the top 100 genes that are significantly co-expressed in specific cell types/intermediate differentiation stages. In all differentiation pathways and for each differentiation stage we found that transcription factors were among the most significant genes distinguishing each cell type (Figure 5A). Moreover, ontology analysis (Ashburner et al. 2000; The Gene Ontology Consortium 2015) using the different subsets of genes revealed strong enrichment of genes regulating the specification of each cell type (Supplemental Table S3).

Since we found that: a) sub-network communities associated with transcription factor activity were identified in all RT correlated networks (Figure 2); b) interactions between TFs in TRNs significantly overlap with RT networks (Figure 4); and c) TF expression patterns distinguish each cell type (Figure 5A); we hypothesize that gene regulation by TFs might be critical not only for cell type-specific transcriptional program establishment of but also for RT program control during cell fate specification. To analyze the potential role of TF in RT regulation we constructed bipartite networks in which we identified the co-regulated patterns of RT that correlated with the expression levels of the TRNs. First, we identified the genes whose RT patterns are correlated with the expression levels of the top 100 genes that distinguish each cell type. Exemplary gene expression levels for a subset of TFs critical for pancreas and liver differentiation are shown in Figure 5B-C, as well as the corresponding genes with correlated patterns for RT regulation. Then, we constructed bipartite networks that consist of two independent but interconnected networks: the TRN side contains genes that are co-expressed in specific developmental stages/cell types and the RT side contains genes whose RT changes were highly correlated with the expression patterns from the TRN side. Importantly, these gene interactions are ***<u>in trans</u>*** and cannot be explained by the “simple” correlation between RT and transcription but correlated transcriptional changes with ***<u>remote, unlinked</u>*** RT changes. Hence, the regulatory interactions described here cannot be explained by the previously recognized correlation. Interaction edges between each side of the bipartite networks were established for nodes in the RT side that correlate (R≥0.75) with the transcriptional levels of at least 50% of the genes in the TRN side. Bipartite networks for pancreatic and liver differentiation are shown in Figures 5D-E respectively. TFs that regulate these specific differentiation pathways are highlighted in each bipartite network at the TRN side (Figure 5D-E); and known pancreatic-specific and liver-specific downstream genes, were found in for each bipartite network at the RT side (Figure 5D-E). These results support the hypothesis that establishment of TRNs during cell differentiation is required for RT control.

**Figure 5:**
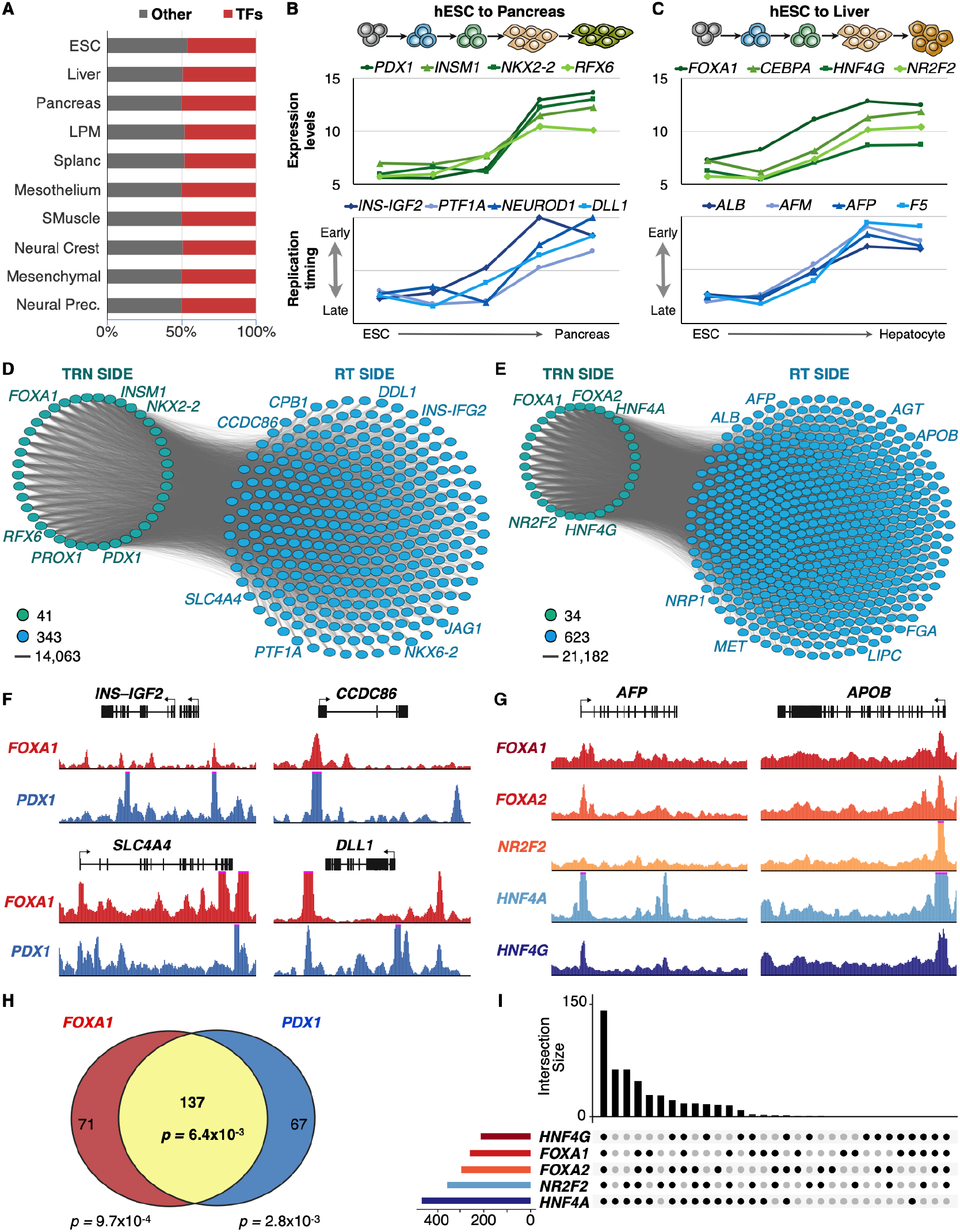
Bipartite networks (click for full resolution). A) Transcription factors expression patterns distinguish each cell type/differentiation intermediate. Cell type-specific expression patterns were analyzed to identify the most significant differentially co-expressed genes, then the resulting genes were classified as TFs or other type. B) Expression patterns of exemplary key TFs of pancreas development correlate with the RT of downstream regulators of pancreatic differentiation. C) Expression patterns of exemplary key TFs of liver development correlate with the RT of downstream regulators of hepatic differentiation. D-E Bipartite networks of pancreas (D) and liver differentiation (E). Bipartite networks were constructed based on the correlation between transcriptional levels of the genes in the TRN side and RT changes of genes in the RT side. TRNs were constructed from the top genes co-expressed in pancreas and liver, respectively. RT network sides were constructed by identification of genes with RT patterns highly correlated with the expression levels of genes at the TRN side. Only edges between TRN and RT networks are shown (with each gene in the RT side connected with at least 50 % of the nodes in the TRN side), as all nodes within each network are interconnected with all other. F) ChIP-seq signals show the co-occupancy of *FOXA1* and *PDX1* at exemplary pancreatic-specific downstream targets predicted by the bipartite network shown in D. G) ChIP-seq signals show the co-occupancy of *FOXA1, FOXA2, NR2F2, HNF4A* and *HNF4G* at hepatic-specific downstream targets predicted by the bipartite networks shown in E. H-I Co-occupancy analysis for the pancreatic-specific and hepatic-specific TFs at the promoters of downstream targets.

During the preparation of this manuscript, ChIP-seq datasets for transcription factors binding became available for liver and pancreas (Diaferia et al. 2016; Wang et al. 2018; https://www.encodeproject.org/). Hence, to further analyze the intriguing relationship of TFs whose expression strongly correlates with the RT of downstream genes *in trans*, we analyzed ChIP-seq data for known TFs required for pancreatic and liver cell fate commitment (Figure 5F-G). For pancreatic differentiation, we analyzed whether downstream targets predicted at the RT side of the bipartite network (Figure 5D), contain binding sites for *FOXA1* (endoderm-specific TF) and *PDX1* (pancreatic-specific TF). We found ChIP-seq peaks for both TFs in close proximity to the promoters (<20kb) of the downstream targets predicted in the bipartite network (Figure 5F). Additionally, we found a highly significant co-occupancy (*p-value*= 6.4×10^-3^) for these TFs at the promoters of predicted targets (Figure 5H). For liver differentiation, we obtained ChIP-seq data for five different TFs required for the regulation of hepatic differentiation (*FOXA1, FOXA2, NR2F2, HNF4A* and *HNF4G*), which allowed us to test for co-occupancy of these key regulators at the predicted targets. Remarkably, we found multiple genes (139) with TF binding peaks at their promoters for all five TFs (Figure 5G and 5I). Moreover, binding of these TFs at the promoters of the predicted targets (Figure 5I) was highly significant for all combinations of TFs co-occupancy when comparing to expected occurrence (Supplemental Table S4). These findings suggest a regulatory link between the establishment of cell type specific TRNs and RT control during lineage specification.

## Discussion

In this study, we introduced a new approach to construct gene regulatory networks exploiting the dynamic changes in DNA replication timing during lineage specification. RT is cell type-specific (Hiratani et al. 2010; Rivera-Mulia et al. 2015; Ryba et al. 2011), regulation of RT is critical to maintain genome stability (Donley et al. 2013; Neelsen et al. 2013; Alver et al. 2014) and abnormal RT is observed in disease (Ryba et al. 2012; Gerhardt et al. 2014; Rivera-Mulia et al. 2017; Sasaki et al. 2017; Rivera-Mulia et al. 2018b). RT is closely related to the spatio-temporal organization of the genome with early and late replicating domains segregating to distinct nuclear compartments (Pope et al. 2014; Rivera-Mulia and Gilbert 2016b). Cell fate commitment is accompanied by dynamic changes in RT that are globally coordinated with transcriptional activity (Rivera-Mulia et al. 2015; Rivera-Mulia and Gilbert 2016b). Hence, RT constitutes a functional readout of genome organization that is linked to gene regulation during cell fate commitment. However, despite a significant correlation of early RT with transcriptional activity, RT and transcription cannot predict each other suggesting an indirect relationship. Here, we constructed RT networks based on RT changes across 15 cell types and differentiation intermediates derived from human embryonic stem cells. We identified thousands of genes from different chromosomes that are co-regulated in RT during cell differentiation (Figure 1C) and constructed distinct RT network models based on their dynamic changes. Our results suggest an intimate link regulation of RT by cell type specific TRNs that is not revealed by association with transcription in cis but is uncovered by RT networks.

Directional RT networks were able to identify sets of unlinked genes whose RT changes coordinately during differentiation, the master regulators of RT changes and their temporally downstream targets. To validate our model of directional RT networks we demonstrated that the gene interactions identified by our novel networks could predict the downstream targets of known regulators of specific differentiation pathways for which ChIP-seq data were available (Figure 3). The algorithms to construct these RT networks that we present here can be applied to explore the interactions of any gene of interest (see Methods for detailed information on the computational pipeline).

Combining TRNs and RT networks into composite and bipartite networks revealed new insights into gene regulation during cell fate commitment. First, we found that there is a highly significant overlap between TRNs and RT networks of TFs (Figure 4A). However, the RT network motifs contain more RT edges than transcriptional edges, suggesting that the TF genes whose RT programs are co-regulated during differentiation do not regulate each other’s transcription levels during cell fate commitment. Hence, establishment of RT regulatory networks of specific subsets of TFs might be necessary for the proper regulation (either at the level of RT or gene expression) of downstream genes in a more complex regulatory circuit to ensure and maintain the distinct cell identities. For example, TRNs may regulate RT independent of their direct role in transcriptional regulation, which in turn affects the responsiveness of RT-regulated genes to downstream transcriptional regulation. Second, composite networks solved the conundrum of high overlap between RT and TRNs with a lack of transcriptional interactions within the RT networks motifs; composite networks revealed more complex circuitries in which transcriptional edges connected otherwise separated RT motifs (Figure 4D-E). Finally, an important question in the DNA replication field is what regulates the replication-timing program. Correlations with chromatin features have been described for decades (Rivera-Mulia and Gilbert 2016b), but we do not understand yet how RT is regulated and how it is re-modeled during cell fate specification. Here, we identified an intriguing relationship of TFs whose expression strongly correlates with the RT of downstream genes *in trans* (Figure 5B-C) and constructed a novel class of bipartite networks (Figure 5D-E). Bipartite networks allowed us to identify hundreds of RT co-regulated genes correlated with expression levels of co-expressed TFs within the same differentiation pathways. This is of particular significance to our understanding of RT control because, despite the correlation between early replication and transcriptional activity, no causal links have been unveiled (Rivera-Mulia and Gilbert 2016a; 2016b; Hiratani et al. 2010; Rivera-Mulia et al. 2015; Zhao et al. 2017). In fact, individual knockout/ knockdown or overexpression of many transcription and chromatin structure regulators (including TFs such as C-MYC, N-MYC, MYOD and PAX5) has no effect on RT (Dileep et al. 2015) and combinatorial coregulation of multiple TFs might be required to control TRNs (Gerstein et al. 2012; Novershtern et al. 2011). Hence, establishment of complex circuitries/complete regulatory TFs networks, rather than transcriptional induction of specific downstream targets, might be required to shape the RT program during development. Since the chromatin is assembled at the replication fork and different types of chromatin are assembled at different times during S phase (Lande-Diner et al. 2009), a TRN change stimulating an RT change would alter the chromatin structure of an entire replication domain and all the genes within that domain contributing to the regulation of nuclear function and organization during cell fate commitment. Overall, the RT networks suggest a novel hypothesis that can be tested to unveil the mechanisms for RT regulation during lineage specification. Experimental manipulation of TFs followed by differentiation protocols would provide novel insights of whether TFs regulate RT independently of their transcriptional roles, whether all TFs are able to regulate RT or only a specific class of TFs have this property and if a complex combinatorial co-occupancy of several TFs and/or binding to super-enhancers is required to remodel the RT program.

## Methods

### Extraction of RT values at the TSS of NCBI RefSeq genes

Replication Timing (RT) data from multiple cell types and intermediate differentiation stages derived from human embryonic stem cells (Rivera-Mulia et al., 2015) were used to extract the RT values at the transcription start sites (TSS) of all RefSeqs genes in R. Briefly, average RT profiles were obtained from replicates and TSS positions were used to predict RT values from the loess smoothed RT profiles (Ryba et al, 2011). This data consists of RT values at the TSS of all RefSeq genes from 15 cell types derived from hESC representing three main germ layers; ectoderm, mesoderm, and endoderm (Figure 1A).

### Construction of RT networks based on coordinated changes in RT

To construct correlated RT networks we denoted the set of genes as G = {g_1, g_2, …, g_n} and the set of cell types as {c_1, c_2, …, c_s}. Then three filters were applied to include exclusively genes that change RT and are co-regulated in during differentiation. First, we removed all genes that do not change RT during cell differentiation. Since all genes that have the same (or very similar) RT values across all cell types will yield high correlation values regardless of their RT values, it will lead into false positive correlations. Thus, we removed all RT constitutive genes and included only the genes that change RT during cell differentiation between the very early (>0.3) and very late (<-0.3) replication. It is worth noting that this is an aggressive filter as these parameters would consider some genes as constitutive even if they may have high variation in RT although it is always early or late in replication. Second, we removed gene pairs with co-regulated RT patterns that are located within the same replication domain. If two genes are located close to each other on the same chromosome, their RT values are expected to be highly correlated due to a natural outcome of the DNA replication process. Such correlations have less significance as compared to those among physically distant genes, for the correlations between distant genes provide hints about the existence of complex interactions that regulate the order in which genes are replicated. Since we have demonstrated that replication domains vary in size from 0.4 to 0.8 Mb (Hiratani et al., 2008; Pope et al., 2014; Rivera-Mulia et al., 2015), we removed all gene pairs separated by <500 Kb (distance threshold denoted as μ). Thus, for all possible gene pairs in G we obtained the locations of g_i1 and g_i2 on the DNA. If they are on the same chromosome, we classified them as *co-located*; if they are in different chromosomes we classified them as *not co-located.* If the two genes g_i1 and g_i2 are *co-located*, the distance between their TSS positions d_i1,i2 was calculated and all genes with d_i1,i2 < μ were removed from the analysis. Finally, we established interaction edges exclusively for gene pairs that are significantly correlated. To do so, for each gene g_i ∈ G we constructed a vector w_i with an entry of the replication timing of g_i in cell type c_j. Then, the model of correlated RT network was defined as G = (V, E), where V and E denote the set of nodes and edges respectively. Here ∀i, 1 ≤ i ≤ n, node v_i ∈ V corresponds to gene g_i. For all pairs of genes g_i1, g_i2 ∈ G, we computed the Pearson’s correlation coefficient between their vectors w_i1 and w_i2. Statistical significance for the Pearson’s correlation of each gene pair was calculated as follow:

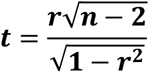

Where the *p-value* being **2 × *P*(*T* > *t*), *r*** is the correlation coefficient and *n* is the number of observations. Next, we calculated the Bonferroni’s corrected *p-values* as:

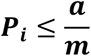

Where ***P_i_*** is the significant Bonferroni-corrected *p-value*, **a** is the significance threshold and ***m*** is the number of tests. Hence, interaction edges between nodes v_i1 and v_i2 were drawn in E if g_i1 and g_i2 were significantly correlated (*p-value* < ***P_i_***).

Correlated RT networks were constructed across all samples, as well as for distinct subsets of cell types focusing on the three major germ layers: ectoderm, mesoderm, and endoderm. In our correlated RT network models, each node represents a gene, and each edge represents a relationship between co-regulated RT of the corresponding two genes in the network.

### RT networks visualization

To visualize our RT network, we focused on the subset of highly connected nodes (> 20-degree count). First, we constructed a two-dimensional map of the RT correlated networks using the force-directed layout algorithm in Cytoscape (Shannon et al., 2003), with the edge length being proportional to the Pearson’s correlation strength (Figure 2). Next, we detected sub-network communities by running Louvain community detection algorithm (Blondel et al., 2008). This method is a heuristic method that is based on modularity optimization. Finally, we used SAFE algorithm to annotate functional attributes for communities. SAFE (Baryshnikova, 2016) is an automated network annotation algorithm. Given a biological network and a set of functional groups or quantitative features of interest, SAFE performs local enrichment analysis to determine which regions of the network are significantly over-represented for each group of features. Thus, local neighborhoods were identified and functional attributes were annotated based on the gene ontology (GO) terms (Baryshnikova 2016).

### Construction of Directional RT networks

Directional RT networks were generated for each differentiation pathway for ‘late replicated to early replicated’ (LtoE) and ‘early replicated to late replicated’ (EtoL). In our late replicated to early replicated (LtoE) directed RT network model each node represents a gene that switches from late replicated to early replicated (LtoE). In creating these networks, we do not consider how much correlated two corresponding genes are. We only consider temporal order of RT changes between two corresponding genes. For differentiation pathway ESCs (the earliest stage) -> Lateral plate mesoderm -> Splanchnic mesoderm -> Smooth muscle (the latest stage) of late replicated to early replicated (LtoE) directed RT network, we draw a directed edge from a gene that switches in earlier stage to a gene that switches in later stage only if the difference of switching stage is one or two. For example, if LtoE pattern of gene g_1 is L->E->E->E in the differentiation pathway ESCs -> Lateral plate mesoderm -> Splanchnic mesoderm -> Smooth muscle and LtoE pattern of gene g_2 is L->L->E->E in the same pathway, we draw an directed edge from g_1 to g_2 as g_1 switches in Lateral plate mesoderm stage (earlier) and g_2 switches in Splanchnic mesoderm stage (later) assuming the change from LtoE of gene g_1 could be causally linked to the change of gene g_2 in the next stage.

### RT network edges overlap with known transcriptional regulatory interactions

We compared the topologies of the RT networks with those of Transcriptional Regulatory Networks (TRNs). Particularly, we used the TRNs constructed using TFs (Neph et al., 2013). To do that, for each cell lineage, we counted the number of edges common to its RT network and TRN. Since interaction edges in a correlated RT networks are not directional, we did not take the direction of edge in TRN into consideration. An undirected edge in the RT network overlaps with a directed edge in the TRN if the gene pairs corresponding to an edge are same in the RT network and the TRN. Using the number of common edges in the two networks, we calculate the *p-value* of the overlap from their hypergeometric distributions. To do so, we denoted the number of nodes (i.e., genes) in the given TRN with n. The total number of possible edges in the complete graphs with *n* nodes is 2×_n_C_2_ (i.e., the number of permutations of two nodes) was denoted as *M*. The number of edges in the TRN was denoted a *m*, the number of edges in the RT network as *K* and the number of common edges between TRN and RT networks with *k*. Randomly generated networks with the same nodes as those in the RT network were constructed and the number of edges common with the given RT was denoted as X. Then, we computed the probability that the number of common edges between the two networks is equal to a given specific value (say *i*) as P (X = i) = (_k_C_i_ × _M-k_C_m-i_) / (_M_C_m_). The numerator in this probability mass function (PMF) describes the number of ways to pick exactly *i* edges from the RT network in *m* draws from a complete graph, without replacement. The denominator shows the number of alternative network topologies with the same nodes as the RT network, which has m edges. Using this PMF, we calculate the *p-value* of having more than or equal to *k* common edges between the RT network and the TRN as 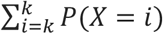. Thus, significant *p-values* identify unexpectedly large number of common edges between the two networks.

### RT network motif identification

Network motifs are defined as recurrent and statistically significant sub-graphs or patterns. We identified the most frequent motifs in the RT networks by creating all possible shapes of connected nodes (in terms of undirected edges) for two, three, and four nodes sub-graphs. The analysis was limited to motifs of ≤ 4 nodes. First, because it has been shown that the fundamental regulatory sub-network patterns consist of a small number of nodes networks (Baiser et al., 2015; Elhesha and Kahveci, 2016; Milo et al., 2002; Yeger-Lotem et al., 2004). Second, because the number of possible motif topologies grow exponentially with number of nodes making it impossible to test larger motif sizes. To identify the most frequent motifs in the RT networks, we counted the number of matching motifs between the RT network and randomized networks. We also created multiple shuffled networks that have the same number of nodes and edges with the RT network and set a *z-score* as 2.54 for a sub-graph to be considered as a motif present in the network. Is important to note that the motif frequency (i.e., the number of times a given motif appears in a given network) do not monotonically decrease or increase with the motif size and the statistical significance of the motif abundance is independent of the motif size and topology (Elhesha and Kahveci, 2016). This is because motif count does not have downward closure property and a motif is considered as abundant in a given network only if its frequency is significantly higher than the number of times the same motif appears in randomized networks (*p-value* <0.01).

### Construction of Composite RT and gene expression networks

The composite network model merges the interactions observed in RT networks (using only genes present in Neph et al., 2013 networks) with the interactions observed in TRNs. To construct the composite networks, we used as base network the motifs (all sub-graphs consisting in connected two, three, and four nodes) detected in the RT networks constructed from the subset of genes present in the TRNs. We then combined the RT and TRNs by taking the union of their edge sets. To do so, we draw the transcriptional edges between the RT nodes according to the interactions identified in the TRNs. Composite networks were visualized in Cytoscape (Shannon et al., 2003).

Transcriptional edges are directional and they were differentiated according to the connectivity within/outside RT motifs, i.e. if a TRN interaction was present within an RT network motif it was visualized as a directed solid edge and it the interaction occurs between nodes from different RT network motifs it was visualized as a directed dashed edge.

### Construction of bipartite RT and transcriptional networks

A bipartite network is a graph with two components. Each component is a set of nodes. In our model, first component is based on the expression patterns of genes, and the second component is based in the replication timing of genes. For our analysis, we started with the list of genes coexpressed in specific cell types that constitute the first component. We append an edge between the first component and the second component if replication timing of a gene in the second component is correlated with expression of a gene in the first component with more than a certain correlation threshold. Next, we removed a gene in the second component if the number of edges of this gene is less than the total number of genes in the first component * ‘ratio’. In this way, we generated the list of genes that constitute the second component.

### ChIP-seq data analysis

ChIP-seq peaks and aligned reads (hg19 genome assembly) for *FOXA1* (ENCSR735KEY), *FOXA2* (ENCSR310NYI), *NR2F2* (ENCSR338MMB), *HNF4A* (ENCSR601OGE), and *HNF4G* (ENCSR297GII) transcription factors in liver tissue were downloaded from ENCODE data portal (https://www.encodeproject.org/). *FOXA1* transcription factor specific peaks and raw reads in pancreas tissue were collected from (Diaferia et al. 2016) (GSE64557), and *PDX1* specific ChIP-seq data in pancreas tissue was downloaded from (Wang et al. 2018) (GSE106949). Aligned reads for *FOXA1* and *PDX1* in the pancreas tissue on hg19 genome assembly were generated using bowtie2 alignment program (Ben Langmead and Salzberg 2012). PDX1 specific peak calling against the respective input was performed using the MACS2 program (Zhang et al. 2008) with parameters: “g hs–q 0.05”. All the significant transcription factor peaks (FDR < 0.05) were retained for downstream analysis, and the raw ChIP-seq signal tracks were scaled to the signal track with minimum coverage for visualization purpose. Individual transcription factor peaks and their respective combinations in liver and pancreas were mapped to the annotated (Harrow et al. 2012) hg19 transcription start sites (+/-20Kb) using “bedtools map” function (Quinlan and Hall 2010) with default parameters. Overlap significance and enrichment of transcription factor binding in RT-network specific genes was measured using Fisher’s exact test by comparing against a similar number of random set of non-RT-network genes.

## Data Access

RT and gene expression datasets were originally published in Rivera-Mulia, et al., 2015 and are publicly available in our laboratory database at http://www.replicationdomain.org, as well as in the ENCODE portal (www.encodeproject.org/). *FOXA1, FOXA2, NR2F2, HNF4A* and *HNF4G* ChIP-seq datasets from liver tissue were obtained from the ENCODE portal (www.encodeproject.org/). *FOXA1* and *PDX1* ChIP-seq datasets from pancreas tissue were obtained from Diaferia et al., 2016 (GSE64557) and Wang et al., 2018 (GSE106949) respectively.

## Acknowledgments

This work was supported by NIH grant GM083337 (D.M.G.).

## Author’s contributions

JCRM, TK and DMG conceived and designed the study; JCRM, SK, HG, AC and FA performed data analysis and interpretation; JCRM and DMG wrote the manuscript.

## Supplemental Material

**Supplemental Table S1.**
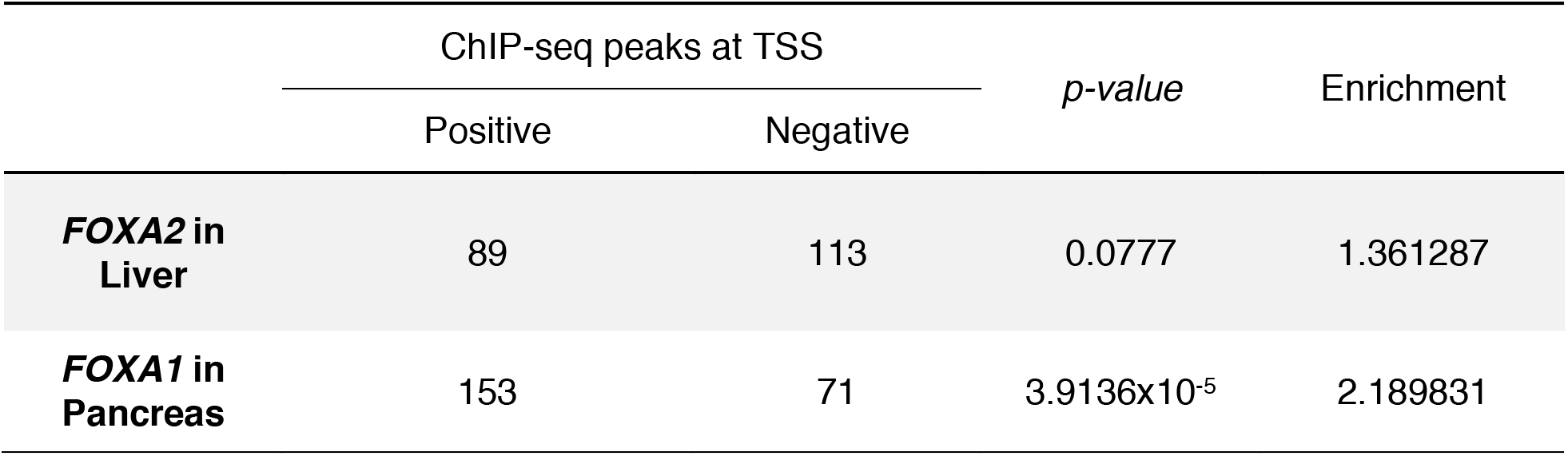
Overlap significance and enrichment of transcription factor binding in directed RT-networks. ChlP-seq data for *FOXA2* transcription factor in liver tissue were downloaded from ENCODE data portal (https://www.encodeproject.org/). ChIP-seq data for *FOXA1* transcription factor in pancreas tissue were collected from Diaferia et al. 2016 (GSE64557). Peak calling against the respective input was performed using the MACS2 program with parameters: “-g hs-q 0.05”. All significant transcription factor peaks (FDR < 0.05) were retained for downstream analysis. Individual transcription factor peaks were mapped to the annotated hg19 transcription start sites (+/-20Kb) using “bedtools map” function (Quinlan and Hall 2010) with default parameters. Overlap significance and enrichment of transcription factor binding in RT-network specific genes was measured using Fisher’s exact test by comparing against a similar number of random set of non-RT-network genes.

|  | ChIP-seq peaks at TSS |  | *p*-value | Enrichment |
| --- | --- | --- | --- | --- |
|  | Positive | Negative |  |  |
| **FOXA2 in Liver** | 89 | 113 | 0.0777 | 1.361287 |
| **FOXA1 in Pancreas** | 153 | 71 | 3.9136x10<sup>-5</sup> | 2.189831 |

**Supplemental Table S2.** Overlap analysis of RT and TRNs interaction edges. RT networks were constructed for matching cell types in the TRNs (Neph, et al., 2013) and common and unique interaction edges were identified. Only genes within the TRNs were used (475 transcription factors). Hypergeometric test was performed to test the overlap significance (*p-values* are shown). Ectoderm cell types = neural crest, mesenchymal stem cells and neural precursor cells. Mesoderm cell types = lateral plate mesoderm, splanchnic mesoderm, mesothelium and smooth muscle. Endoderm = definitive endoderm, immature hepatic, hepatoblast, liver (hepatocytes), primitive gut, posterior foregut and pancreas (pancreatic endoderm).

|  | all<br>differentiation<br>pathways | ectoderm | mesoderm | endoderm |
| --- | --- | --- | --- | --- |
| RT network size | 96 | 853 | 623 | 204 |
| TRN size | 3303 | 843 | 865 | 292 |
| Common edges | 15 | 108 | 71 | 9 |
| RT network-specific edges | 81 | 745 | 552 | 195 |
| TRN-specific edges | 3288 | 735 | 794 | 283 |
| Hypergeometric *p-value* | 0.04502 | 4.619e-12 | 2.815e-07 | 0.32358 |

**Supplemental Table S3.**
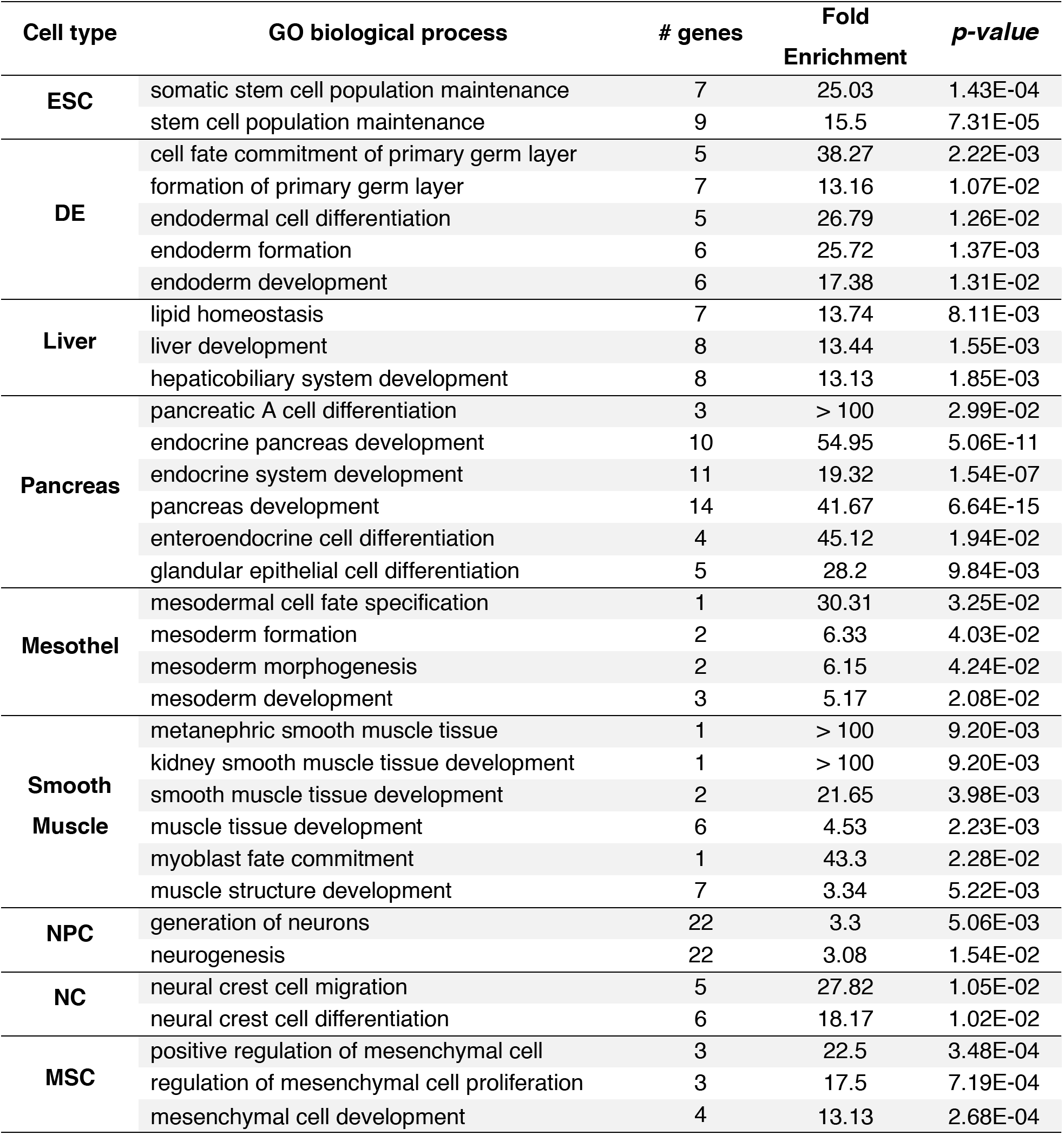
GO analysis of co-expressed genes in each cell type. Co-expressed genes were identified by weighted correlation network analysis (Langfelder and Horvath, 2008) and ontology analysis (Ashburner et al., 2000; The Gene Ontology Consortium, 2015) using the top 100 genes was performed for each cell type.

**Supplemental Table S4.**
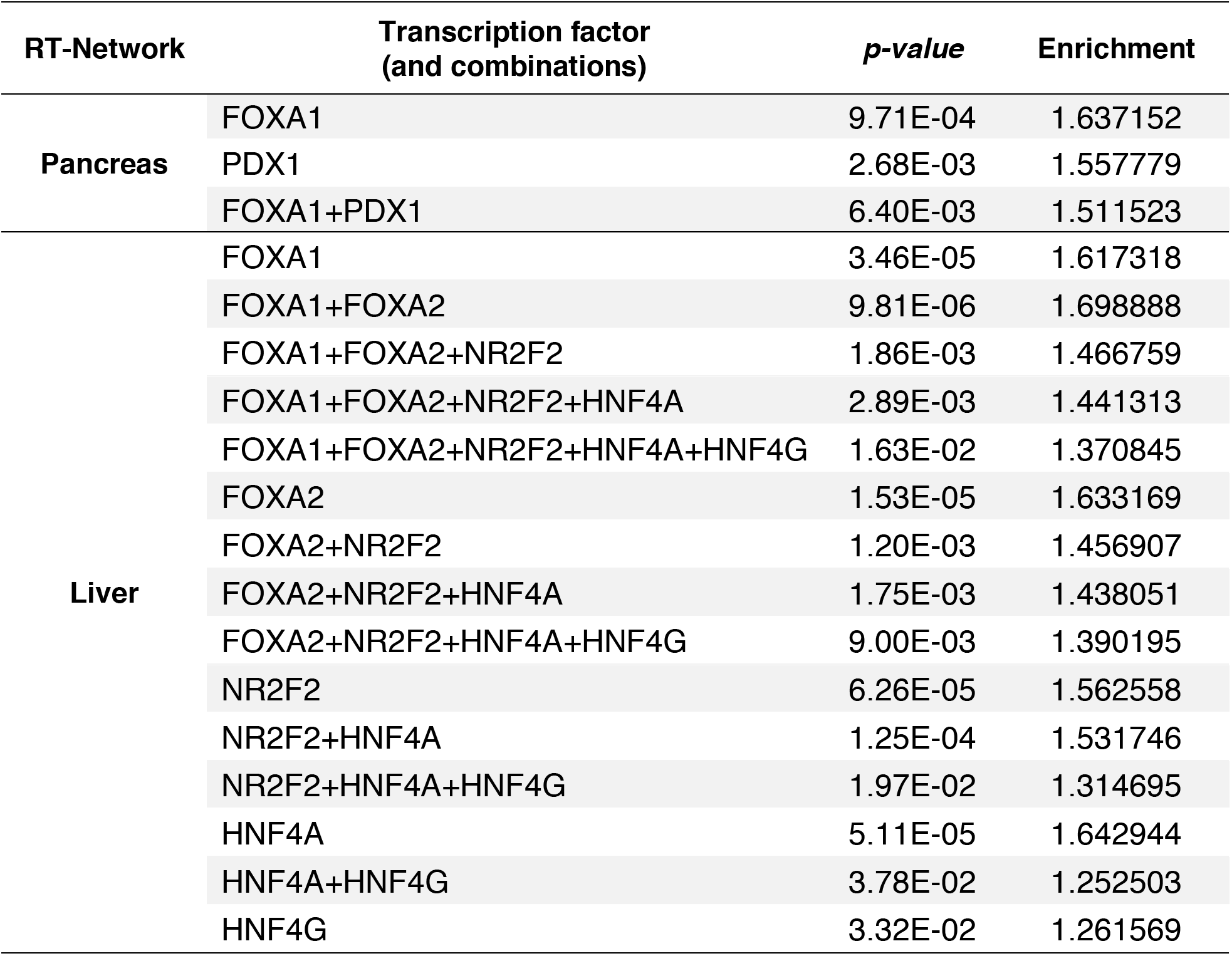
Overlap significance and enrichment of transcription factor binding in bipartite RT-networks. ChIP-seq data for*FOXA1, FOXA2, NR2F2, HNF4A*, and *HNF4G* transcription factors in liver tissue were downloaded from ENCODE data portal (www.encodeproject.org/). ChIP-seq data for *FOXA1* transcription factor in pancreas tissue were collected from Diaferia et al. 2016 (GSE64557) and PDX1 specific ChIP-seq data in pancreas tissue was downloaded from Wang et al. 2018 (GSE106949). Aligned reads on hg19 genome assembly were generated using bowtie2 alignment program. Specific peak calling against respective input was performed using the MACS2 program. All significant transcription factor peaks (FDR < 0.05) were retained for downstream analysis. Individual transcription factor peaks were mapped to the annotated hg19 transcription start sites (+/-20Kb) using “bedtools map” function with default parameters. Overlap significance and enrichment of transcription factor binding in RT-network specific genes was measured using Fisher’s exact test by comparing against a similar number of random set of non-RT-network genes.

**Supplemental Figure S1.**
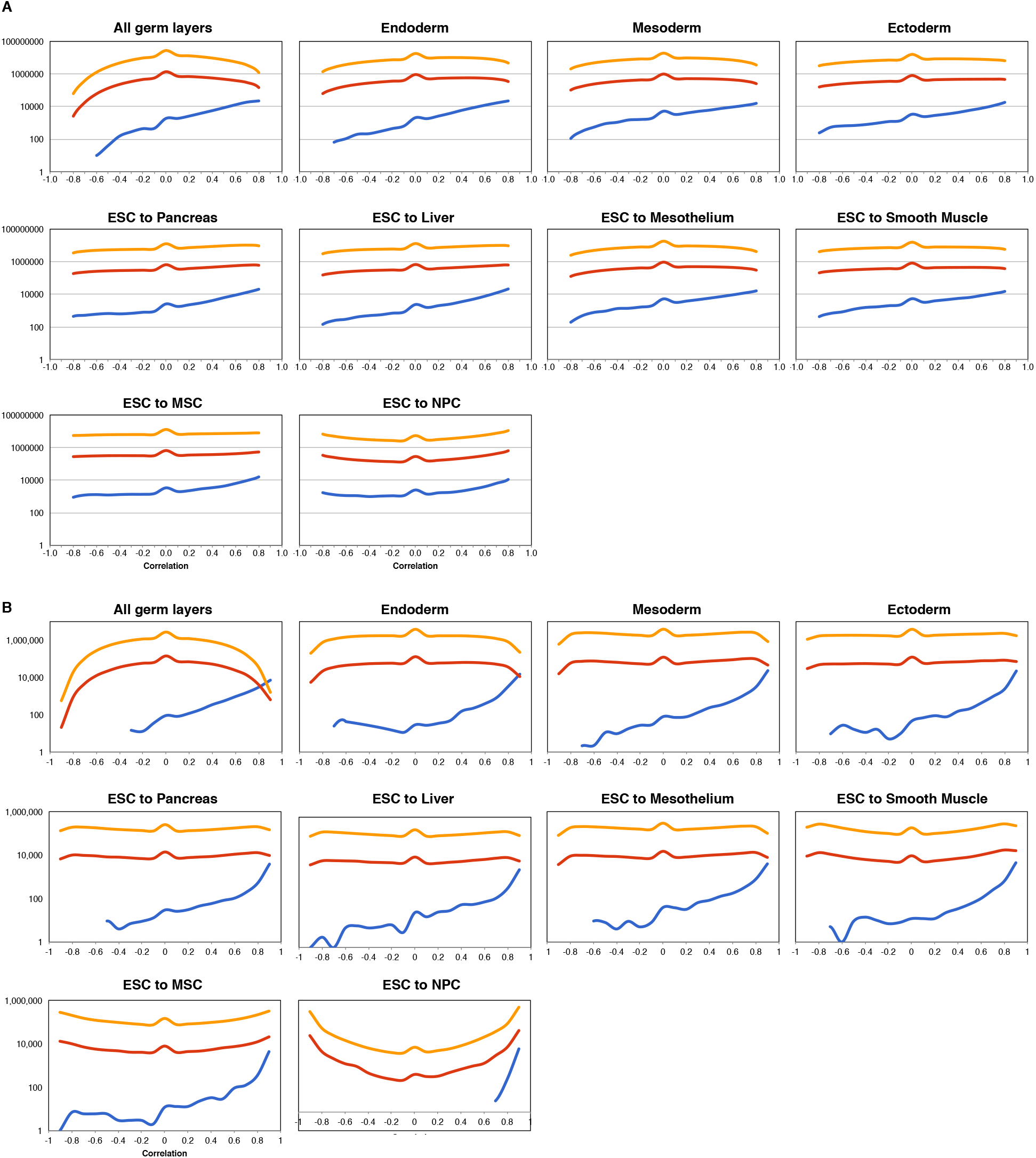
RT correlation per gene pairs. Number of gene pairs as function of RT correlation for distinct categories of gene pairs: co-located close (within 500kb), co-located distant (separated by > 500kb) and not co-located (from different chromosomes). All gene pairs were computed in (A) and gene pairs between genes that change RT significantly within each differentiation pathway (B) are shown.

**Supplemental Figure S2.**
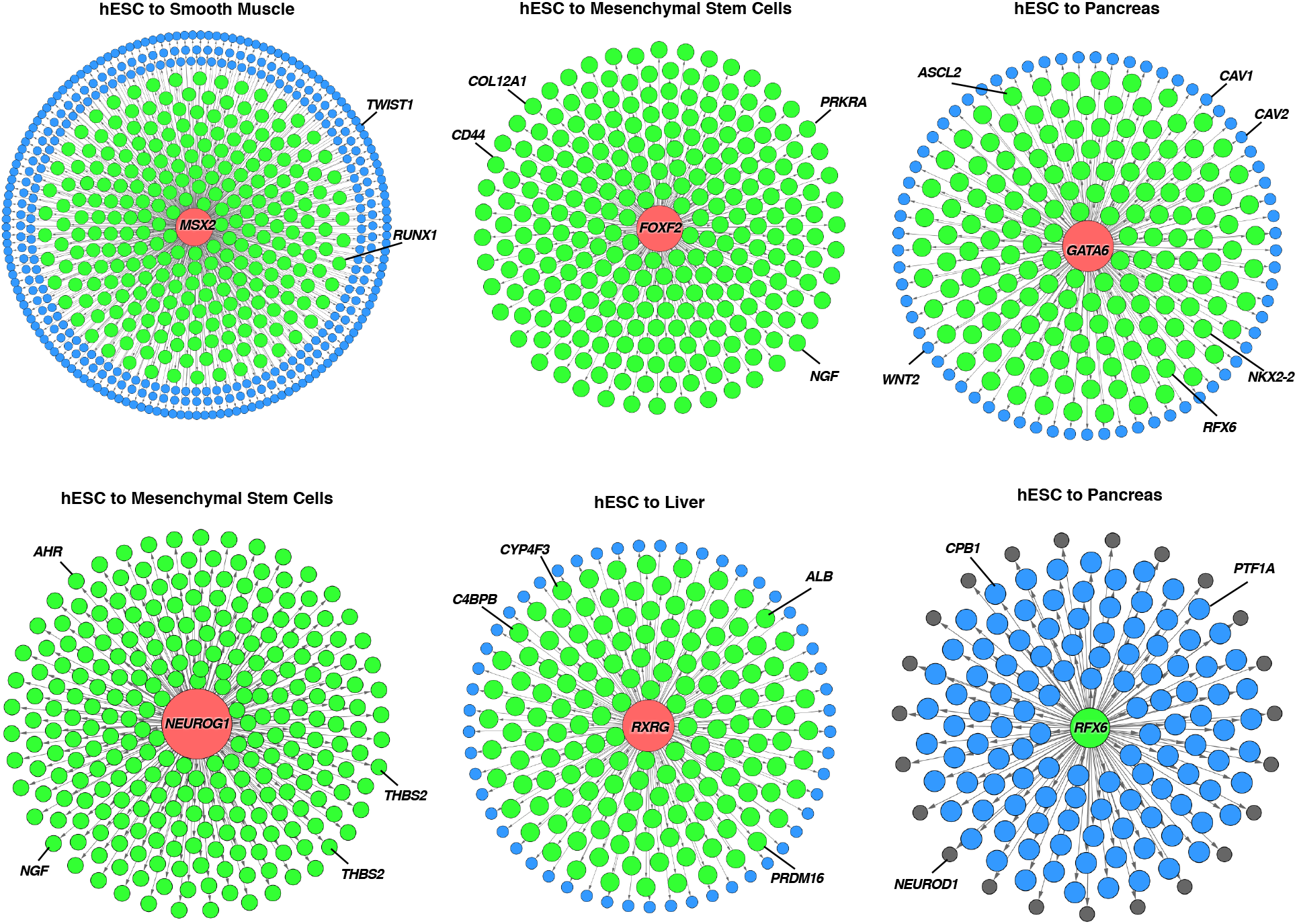
Directional RT networks. Exemplary directional RT networks for known key regulators of each differentiation pathway are shown. Genes that change RT during cell differentiation were classified according to their temporal order of RT changes: *“master regulators*” were defined as the genes that change RT in the earliest differentiation transition and have the largest degree of connectivity (red nodes), while downstream nodes were classified according to the time during differentiation when they change RT as *managers* or *effectors* (green and blue nodes and grey nodes respectively). Known TFs involved in regulation cell differentiation for each pathway were selected as source nodes and potential downstream targets were identified based on the temporal RT changes. 2D maps of directional RT networks were then visualized in in Cytoscape (Shannon et al., 2003). Known cell type-specific genes that distinguish each lineage were found among the downstream genes in directed RT networks (exemplary genes are highlighted in each directed RT network).

**Supplemental Figure S3.**
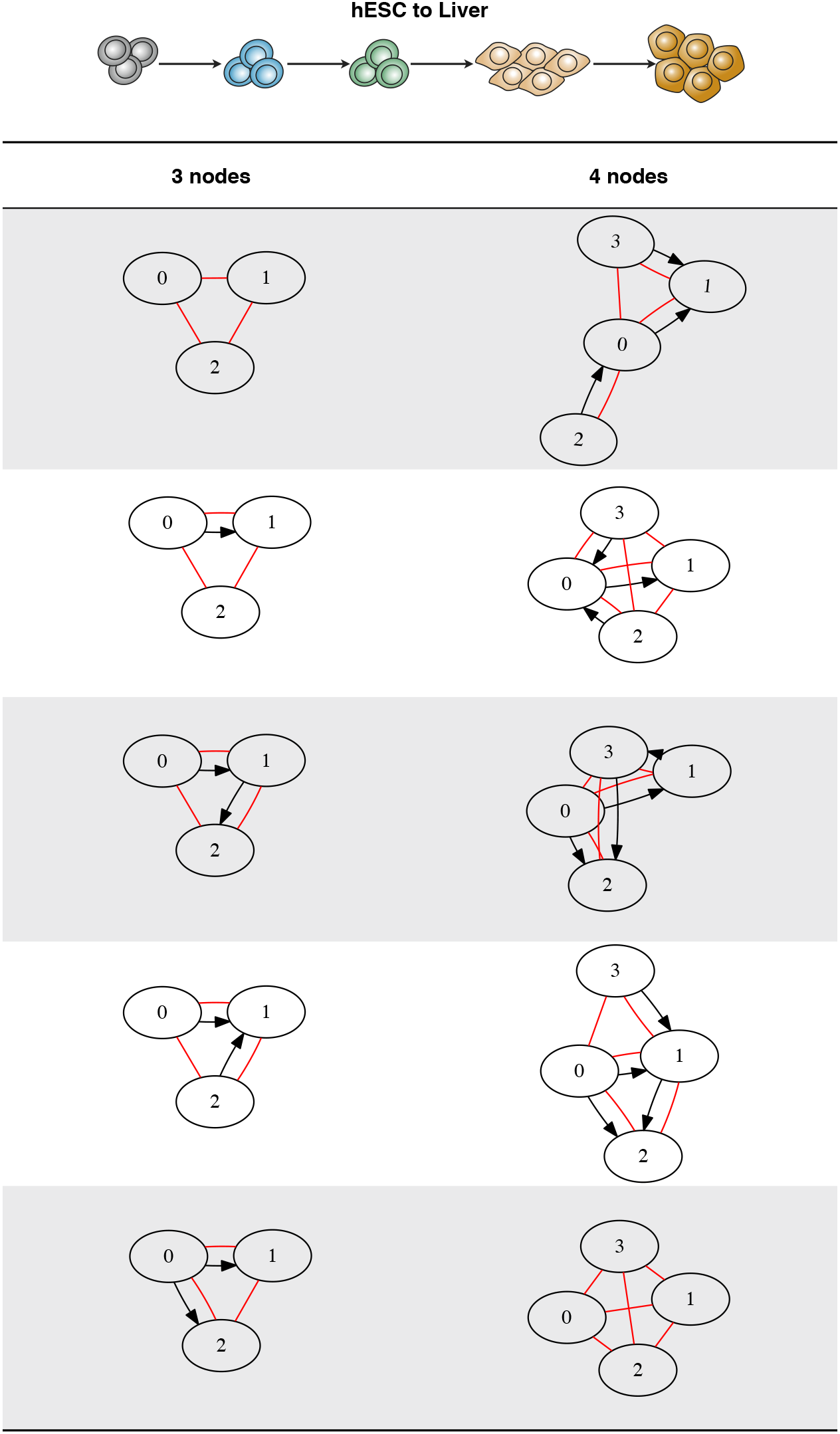
Top 5 motifs identified in liver RT networks. All possible motifs composed were computed and the most enriched motifs were identified. Statistical significance of each motif pattern was calculated by comparison to randomized networks (Baiser et al., 2015; Elhesha and Kahveci, 2016; Milo et al., 2002). Shown are the 5 most enriched motifs in liver RT networks. RT edges are shorn in red (undirected edges) and TRN edges are shown in black (directed edges).

**Supplemental Figure S4.**
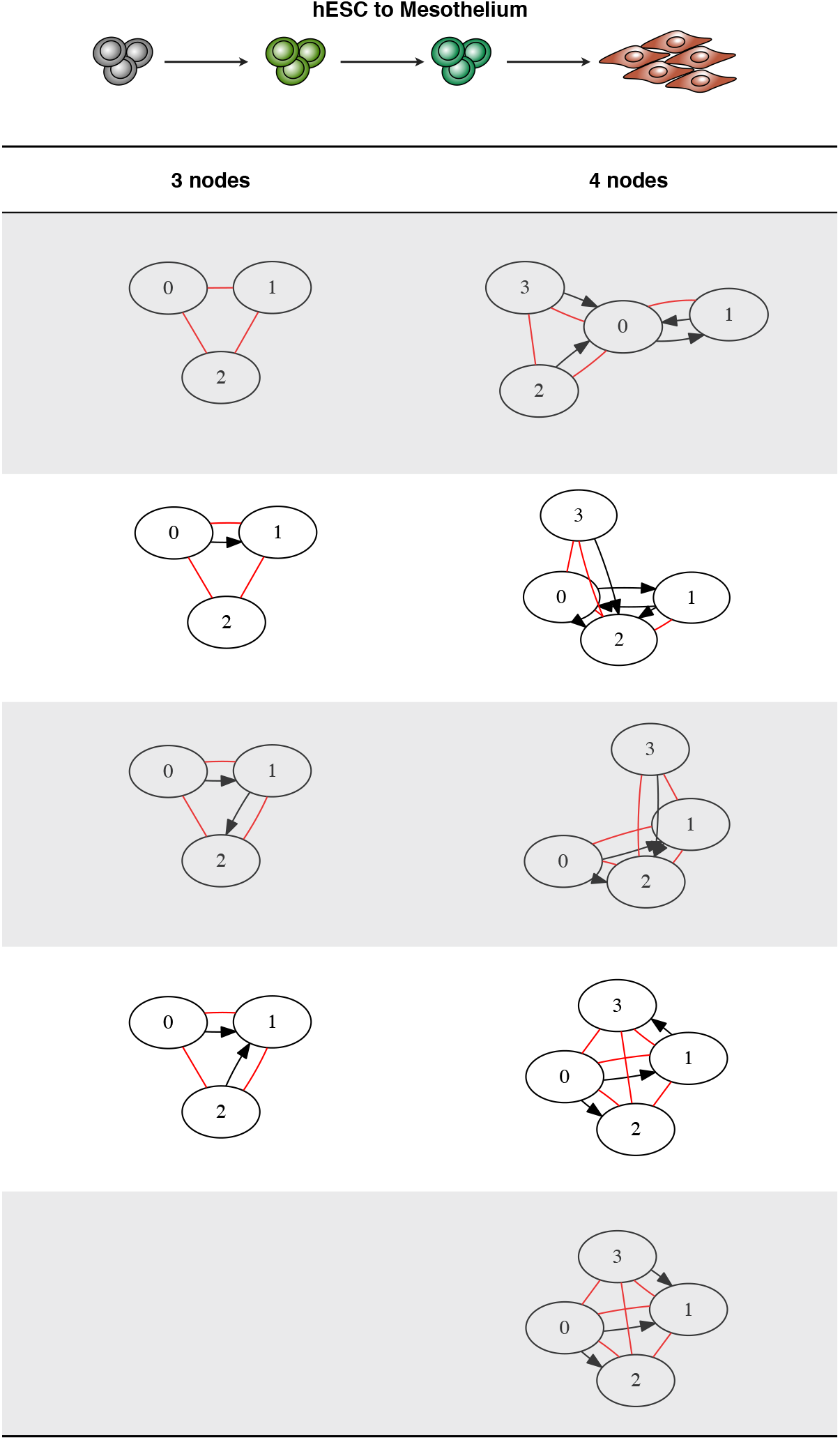
Top 5 motifs identified in mesothelium RT networks. All possible motifs composed by 2-4 nodes were computed and the most enriched motifs were identified. Statistical significance of each motif pattern was calculated by comparison to randomized networks (Baiser et al., 2015; Elhesha and Kahveci, 2016; Milo et al., 2002). Shown are the 5 most enriched motifs in mesothelium RT networks. RT edges are shorn in red (undirected edges) and TRN edges are shown in black (directed edges).

**Supplemental Figure S5.**
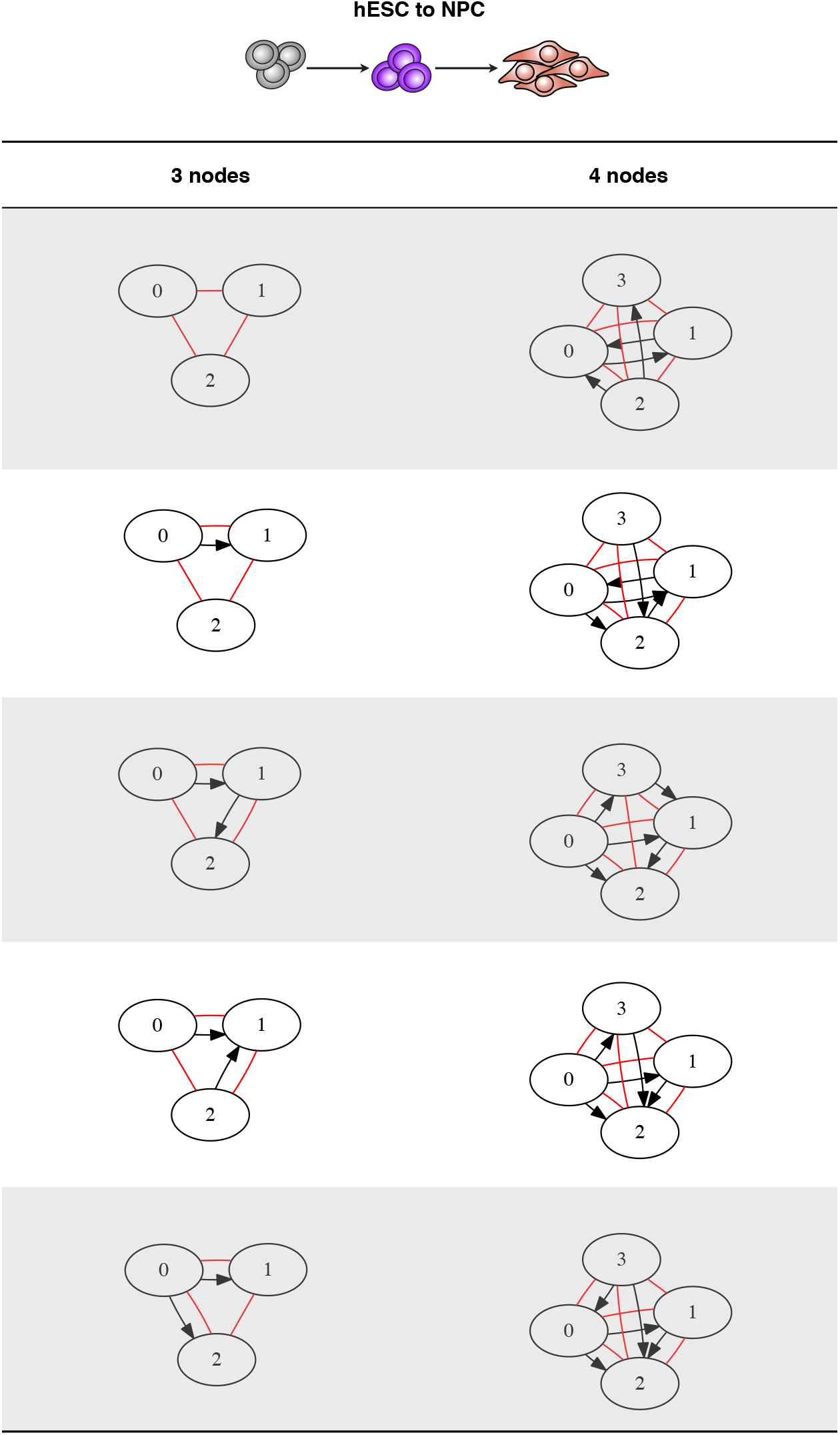
Top 5 motifs identified in NPC RT networks. All possible motifs composed by 2-4 nodes were computed and the most enriched motifs were identified. Statistical significance of each motif pattern was calculated by comparison to randomized networks (Baiser et al., 2015; Elhesha and Kahveci, 2016; Milo et al., 2002). Shown are the 5 most enriched motifs in NPC RT networks. RT edges are shorn in red (undirected edges) and TRN edges are shown in black (directed edges).

